# Conserved pheromone production, response and degradation by *Streptococcus mutans*

**DOI:** 10.1101/635508

**Authors:** Antonio Pedro Ricomini Filho, Rabia Khan, Heidi Aarø Åmdal, Fernanda C. Petersen

## Abstract

*Streptococcus mutans*, a bacterium with high cariogenic potential, coordinates competence for natural transformation and bacteriocin production via the XIP and CSP pheromones. CSP is effective in inducing bacteriocin responses, but not competence in chemically defined media (CDM). This is in contrast to XIP, which is a strong inducer of competence in CDM, but can also stimulate bacteriocin genes as a late response. Inter-connections between the pathways activated by the two pheromones have been characterized in certain detail in *S. mutans* UA159, but it is mostly unknown whether such findings are representative for the species. In this study, we used bioassays based on luciferase reporters for the bacteriocin gene *cipB* and the alternative sigma factor *sigX* to investigate various *S. mutans* isolates for production and response to CSP and XIP pheromones in CDM. Similar to *S. mutans* UA159, endogenous CSP was undetectable in the culture supernatants of all tested strains. During optimization of the bioassay using the *cipB* reporter, we discovered that the acivity of exogenous CSP used as a standard was reduced over time during *S. mutans* growth. Using a FRET-CSP reporter peptide, we found that *S. mutans* UA159 was indeed able to degrade CSP, and that such proteolytic activity was not significantly different in isogenic mutants with deletion of the protease gene *htrA*, or the competence genes *sigX, oppD*, and *comR*. CSP proteolysis was also detected in all the wild type strains, indicating that such activity is conserved in *S. mutans*. For the XIP pheromone, endogenous production was observed in the supernatants of all 34 tested strains at peak concentrations in culture supernatants that varied between 200 nM and 26000 nM. Transformation in the presence of exogenous XIP was detected in all, but one, of the isolates. The efficiency of transformation varied, however, among the different strains, and for those with the highest transformation rates, endogenous XIP peak concentrations in the supernatants were above 2000 nM XIP. We conclude that XIP production and inducing effect on transformation, as well as proteolytic activity leading to the inactivation of CSP are conserved functions among different *S. mutans* isolates. Understanding the functionality and conservation of pheromone systems in *S. mutans* may lead to novel strategies to prevent or treat unbalances in oral microbiomes that may favour diseases.

## INTRODUCTION

Natural genetic transformation is widely distributed in bacteria. In streptococci and in bacillus it occurs during a genetically programmed differentiated state called competence. During this state the bacteria become capable to take up DNA from the environment and incorporate it into their genomes. The capacity for natural transformation has been reported for more than 80 bacterial species (1, 2). Among streptococci, competence for natural transformation includes most species of the mitis, salivarius, bovis, anginosus and mutans groups (1, 3–5). The core of the machinery necessary for streptococcal transformation relies on the transcription of the conserved alternative sigma factor SigX, also known as ComX. SigX orchestrates a core response in streptococcal species characterized by the induction of 27 to 30 genes (6). The functions of the core genes are predominantly related to transformation, most of them coding for competence effector proteins for DNA binding, uptake and recombination (6–8).

*Streptococcus mutans* is a member of the mutans group and part of the human oral microbiota. Environmental factors disturbing the ecological balance in the oral cavity, such as increased sugar intake, may favor *S. mutans* growth. The *S. mutans* acidogenic and aciduric properties may then contribute to tooth demineralization and dental caries (9). Natural transformation was first reported in *S. mutans* in 1981 (10). This and later studies showed that transformation in *S. mutans*, at least in the laboratory setting, is restricted to a limited range of strains, and depends on environmental conditions not yet fully understood (10). Studies investigating the *S. mutans* pan-genome have recently revealed the ubiquity of *sigX* and competence effector genes in *S. mutans* (11–13), indicating that competence may be a conserved feature in *S. mutans*. Moreover, the extensive horizontal gene transfer observed in the genomes of *S. mutans* clinical isolates (11), indicates that transformation occurs in their natural habitat, and may be a widespread feature in the species.

In *S. mutans* the competent state is triggered by two linear peptides, the CSP (<u>c</u>ompetence <u>s</u>timulating <u>p</u>eptide) (7) and the XIP (*sig<u>X</u>*-<u>i</u>nducing <u>p</u>eptide) (Figure 1) (8). CSP was the first described pheromone and, until recently, the only one known to activate the competence system for genetic transformation in *S. mutans*. The *comC* gene, which encodes CSP, is downstream the *comDE* operon, encoding the ComD histidine kinase and the ComE response regulator. At least in the synthetic form, CSP is thought to bind to the ComD histidine kinase of the ComDE two-component system, leading to phosphorylation of the ComE response regulator (14). Phosphorylated ComE directly up-regulates the transcription of clusters of bacteriocin-related genes (6, 8, 15, 16). These include among others the SMU_1914 gene (also known as *nlmC* and *cipB*), encoding mutacin V, and the putative immunity protein SMU_1913, found in a *comE* downstream region. Mutacin V has been proposed to link the early CSP bacteriocin-inducing response to the competence response (17–21). Notably, the encoding gene SMU_1914 is absent in a range of *S. mutans* strains (13), and other genetic elements, perhaps more conserved, may link the two processes. Essential for competence development is the activation of the XIP pheromone encoding gene, *comS*, followed by the upregulation of the alternative sigma factor SigX, the master core regulator of competence. In rich media, ComS seems to be processed to XIP inside the cells. Intracellular XIP and/or ComS then binds to ComR to activate the competence pathway (22). The predicted CSP is in general conserved among *S. mutans* strains (7, 16), and there are indications that *S. mutans* may form a single CSP pherotype (16, 23). At least in complex medium, the CSP precursor peptide is thought to be processed during and after export, resulting in the active CSP (16, 23).

**Figure 1.**
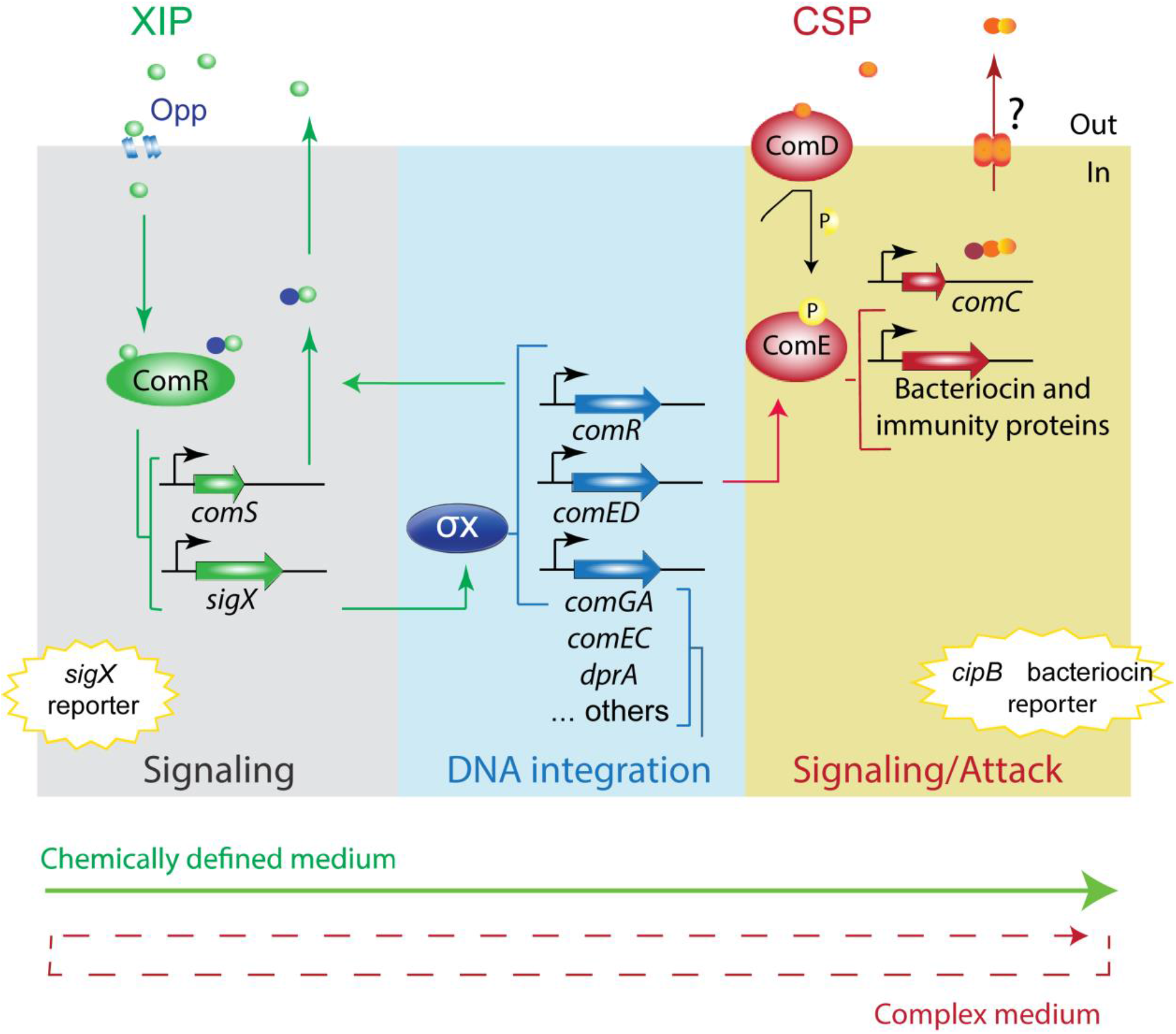
Model of *S. mutans* XIP and CSP pheromone signaling pathways. The model is based on the UA159 *S. mutans* strain. In peptide-free chemically defined media, the *S. mutans* XIP pheromone outside the cells is internalized via the Opp permease, and binds to the ComR regulator to activate *comS*, thus creating a positive feedback loop. ComS is processed to XIP, but can also function in its unprocessed form as an internal signal, to activate the ComR-pheromone signaling pathway. The ComR-pheromone complex also activates *sigX*, coding for the alternative sigma factor X (σX). σX is the master regulator of competence for natural transformation that regulates the expression of the streptococcal core set of genes involved in the competence response, most of them with functions that enable DNA acquisition and integration into its genome. σX induces also *comR* and *comED*. ComED is a two-component system comprising the histidine kinase ComD and the cognate response regulator ComE. ComD autophosphorylates upon CSP pheromone binding, and transfers the phosphoryl group to ComE, which in turn induces the expression of bacteriocin and immunity proteins, including the gene for the non-lantibiotic bacteriocin *cipB*. In rich media, ComE is also thought to induce *comC*, which product is pre-processed and exported by an ABC-exporter, and cleaved to its active form outside the cells (18-CSP pheromone). In CDM, CSP induces the ComED pathway, but it is unknwon whether the CSP pheromone system is endogenously activated. It is also unknown why in complex media the CSP pheromone induces competence, but in CDM, only the bacteriocin pathway is stimulated. To interrogate the conservation of the ComRS and ComED signaling pathways in other members of the species, *sigX* and *cipB* luciferase reporter bioassays were employed.

The XIP pheromone system is found in several streptococci (3, 5, 8, 24, 25). In *S. mutans* UA159 the XIP pheromone encoded by *comS* has been identified in culture supernatants grown in chemically defined medium (CDM) lacking peptides (8, 26). XIP is produced as a pro-peptide (ComS) that is possibly exported and processed into the active XIP (N-GLDWWSL) (4, 8, 26). The presence of an exporter has, however, never been demonstrated, and recent studies inidicate that release of XIP or ComS by autolysis is sufficient to promote intercellular communication in CDM (27). Under such conditions, response to the XIP depends on the Opp oligopeptide permease system, indicating that the peptide is internalized (8). Once inside the cells XIP is thought to bind to the Rgg-like regulator ComR to activate transcription of *sigX* and *comS*. In contrast with XIP, CSP shows no activity or low potency in triggering competence in CDM (28–30), but can still induce the expression of bacteriocin-related genes of the ComE regulon (28). When it comes to transformation, the use of synthetic CSP has led to higher levels of S. mutans transformation (31, 32), but has not extended the range of strains transformed in the absence of the synthetic pheromone. As for the synthetic XIP, evidence for competence induction and endogenous pheromone production has so far been restricted to the reference strain UA159, with few exceptions (12).

In this study we investigated the functional conservation of the *S. mutans* CSP and XIP pheromone signaling systems in CDM. Extracellular CSP activity was not detected in any of the tested strains. Exposure to synthetic CSP revealed that *S. mutans* can indeed degrade CSP, a behavior that was conserved in the examined strains, and that at least in strain UA159 was not abolished by deletion of *sigX, comR, oppD*, or the *htrA* protease gene. For the XIP pheromone, endogenous production was found in all tested strains, and transformation was found in all, but one of the isolates grown in CDM. The results thus indicate the presence of a single *S. mutans* XIP pherogroup, and suggest that *S. mutans* has a conserved ability to suppress CSP activity in CDM.

## MATERIAL AND METHODS

### Bacterial strains and media

The *Streptococcus mutans* strains used in this study and their relevant characteristics are listed in Table 1. The strains used in the study were selected from the collection of strains in our laboratory, representing strains known to be transformed (UA159, V403, OMZ175, LML-2, LML-4, GS5, NG8, BM71, LT11) and others in which genetic competence has not yet been characterized. Todd-Hewitt broth (THB; Becton Dickinson) was used to grow all the strains used for DNA extraction and DNA sequencing. Chemically Defined Medium (CDM) (8) was used to perform the genetic transformation assays and bioassay to measure the activity of exogenously added CSP in the supernatants of *S. mutans*, and also to verify and quantify the concentration of extracellular native XIP in culture supernatants (4, 26). The antibiotics erythromycin, kanamycin and spectinomycin used for mutant construction were used at final concentrations of 10 μg mL^−1^, 500 μg mL^−1^ and 500 μg mL^−1^, respectively. THB agar and THB agar supplemented with erythromycin were used to enumerate the total and transformed number of *S. mutans*.

**Table 1.**
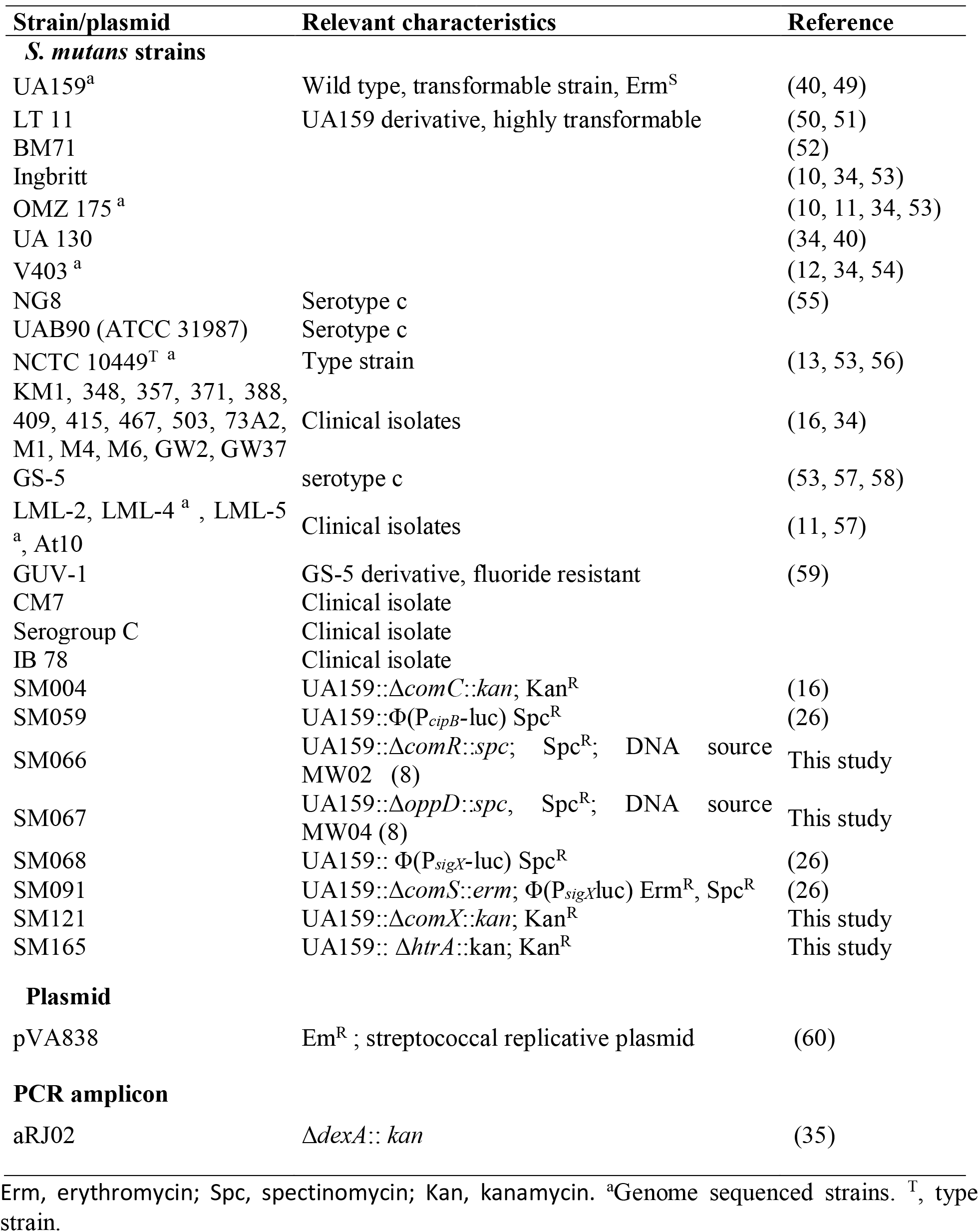
Strains and plasmid used in this study.

#### Construction of mutants

The deletion mutants were constructed using the PCR-ligation mutagenesis strategy (33). Sequence information was obtained from the *S. mutans* UA159 genome. Mutants with deletion of *comX* and *htrA* were constructed. *Asc*I or *Fse*I sites were incorporated into the 5’-ends of the oligonucleotide primers and both ends of the resistance cassette. The kanamycin resistance cassette was amplified using primer pair FP001-FP068. The *comX* flanking regions were amplified with the primers pairs FP462 (CTTGGTAGCAGGAGAGCAC), FP463 (AAAGCACAGCCTGCTTCAAT) and FP714 (TGCCGAACACAGCAGTTAAG), FP715 (CATTCCCTCTTGTTGCCAAT). The *htrA* flanking regions were amplified with the primers pairs FP807 (TCCCTCCAATAACGAAGGTCA), FP808 (GGTAAGTGTTGA TATGACCCCT) and FP809 (GAAGGTAGCGTCTATCAGCGA), FP810 (GCAGTCGAGGTTGATAGGGA). The resultant amplicons were digested with *Asc*I or *Fse*I whereas the kanamycin resistance cassette was digested with both enzymes. The upstream and downstream amplicons of target genes were ligated to the kanamycin cassette using T4 DNA ligase. The two ligated products were mixed and PCR amplified with distal primers. The resultant amplicons were used to transform *S. mutans*. Gene deletion was confirmed by PCR amplification and gel electrophoresis.

### DNA sequence analysis

The chromosomal DNA of *S. mutans* strains were isolated as previously described (34). The primers FP678 (5’-ATGCGGAAGCTAAAAAGAGC-3’) and FP679 (5’-TCCAGTCTTCCTATCTGAGCAA-3’) were used to amplify a region of 431 bp, which contains the tRNA transcriptional terminator of *comR* (9 bp stem, a 4 nt loop and a T-rich region at the 3’ side of the stem-loop), the *comS* promoter and the *comS* sequence. First, a PCR was performed (20 μL reaction volume containing 2.5 mM MgCl_2_, 2 pmol of each primer, 0.2 mM of each dNTP, 2 μL of 10x buffer, 50-100 ng of genomic DNA and 2 units of TaqDNA polymerase using a thermocycling profile of initial denaturing period of 3 minutes (min) at 94°C; followed by 30 cycles of 30 seconds (sec) at 94°C, 30 sec at 55°C and 2.5 min at 72°C; with a final extension period of 3 min at 72°C and the products were visualized on a 1% agarose gel to check the amplified fragments by the primers (34). The sequencing was performed bi-directionally in a single experiment for all the samples using BigDye^®^ Terminator v1.1 Cycle Sequencing Kit (Applied Biosystems, CA, USA) on an ABI 3730 DNA analyzer (Applied Biosystems, CA, USA). All the sequence analysis was done using the Sequencher 5.0 software (Gene Codes, MI, USA). The electropherograms were checked and comparative analysis of the bidirectional sequences was performed using the *S. mutans* UA159 sequence as a reference.

### Bioassay of CSP and XIP activity in culture supernatants

The CSP bioassay was performed to measure the production of native CSP and activity of exogenously added CSP in the supernatants of *S. mutans* during growth, and the XIP bioassay was to verify and quantify the concentration of extracellular native XIP in culture supernatants. A P_*cipB*_-luc reporter (SM059) was used in the CSP bioassay as previously described (26), and a P_*sigX*_-luc reporter in a *comS* deletion background (SM091) was used to perform the XIP bioassay as described by Desai et al. (4), with slight modifications (26).

To estimate CSP production and response, the supernatants were collected by 10x dilution of precultures at OD_600_ of 0,6 in CDM containing 10% Bovine serum albumin (BSA). When added, 250nM CSP concentration was used. Supernatants were collected for 6 hours during growth. For XIP production and response, overnight cultures of *S. mutans* strains were diluted in fresh CDM to an optical density at 600 nm (OD_600_) of 0.05 and incubated in air at 37°C. At each hour, from the second until the tenth hour of bacterial growth, the OD_600_ was measured and 1 mL of the culture was centrifuged (10,000 g for 10 min at 4°C) to collect the supernatant. The indicator strains SM059 and SM091 were grown in CDM to an OD_600_ of 0.05 and stored in microcentrifuge tubes with 10% of glycerol at −80°C. The entire assay was performed with stock cultures from the same batch. The bioassay was performed adding 10 μL of each culture supernatant to 50 μL of the indicator strain, 40 μL fresh CDM and 20 μL of 1.0 mM D-luciferin (Synchem, Felsberg-Altenberg, Germany) in flat-bottom 96-well plate (Nunc, Rochester, EUA). Luminescence was measured by reading the plates in a multidetection microplate reader (Synergy HT; BioTek Instruments, Vermont, USA). To quantify the XIP concentrations in the culture supernatants, standard curves were performed at the same time, replacing the culture supernatant by 10 μL of CDM-standard solutions with different XIP concentrations. CDM without synthetic peptides was used to obtain background values that were subtracted from the sample values. For strains exhibiting no XIP activity in the supernatants, the experiment was repeated at least twice. The software SigmaPlot (version 12.0, Systat Software, Inc.) was used to calculate the XIP concentrations using the relative light unit (RLU) values.

### Genetic transformation

Overnight cultures of all *S. mutans* strains were diluted 1:10 in fresh CDM and incubated in 5% CO_2_ at 37°C. The OD_600_ was followed until the cultures reached an absorbance value of 0.6, then 10% of glycerol was added and they were stored at −80°C. To perform the assay the stored cultures were diluted in fresh CDM 1:10 to an OD_600_ of 0.05 and incubated in air for 2h at 37°C. The OD was measured and aliquots of 150 μL of the bacterial suspensions were transferred to 1.5 mL microcentrifuge tubes. The plasmid pVA838 (Table) was added to all tubes at a final concentration of 1 μg mL^−1^ and XIP was added in the experimental group at a final concentration of 1,000 nM. The samples were then incubated in air for additional 4 h at 37°C, at which time the OD was measured again and the bacterial suspensions were serially diluted (up to 1 to 10^6^) in PBS. The suspensions were plated in duplicate on THB agar and THB agar supplemented with erythromycin 10 μg mL^−1^. The plates were incubated in 5% CO_2_ for 48 h at 37°C before counting the colony-forming units (CFU). The number of genetically transformed cells was divided per total CFU to obtain the values for transformation frequency. Three independent experiments were performed to evaluate transformation in CDM.

In order to evaluate genetic transformation in the non-responsive strains, an independent experiment extending the incubation time to 10 h before plating was performed. In addition, at 6 h an extra load of pVA838 was added, which increased the final concentration to 2 μg mL^−1^. For the remaining non-responsive strains, transformation was attempted using the 6.3kb aRJ02 chromosomal PCR amplicon with a kanamycin marker, which is expected to have a higher sensitivity for detection of transformability (Table) (35, 36).

### Synthetic pheromones

The synthetic XIP (ComS_11-17_; GLDWWSL) (8, 26) and the 18-CSP (ComC_26-43_; SGSLSTFFRLFNRSFTQA) (16) were synthesized by GenScript (GenScript Corporation, NJ, USA), both with an estimated purity of 98% (26). The stock solutions of both peptides were stored in small aliquots at −20°C. The final concentration used in the genetic transformation assays for XIP was 1,000 nM. For FRET-assays to investigate protease activity, the 18-CSP was synthesized using methoxy-coumarin-acetic-acidyl (MCA) on the N-terminal and Lys-Dinitrophenyl on the C-terminal (Dnp) (MCA-SGSLSTFFRLFNRSFTQA-Dnp; GenScript).

### FRET-18CSP peptide cleavage reporter assays

#### Growth in the presence of FRET-18CSP

Pre-cultures of *S. mutans* were diluted in 2 mL CDM supplemented with 2% BSA to an OD_600_ of 0,04 to 0,05, and incubated at 37°C in air atmosphere to an OD_600_ of 0,06. The cultures were then distributed into the wells of a 96-well plate (`115μl in each well`), and 5 μl of `FRET`-18CSP peptide was `added` (`4μM` final concentration). The plate was incubated at 37°C in air, and fluorescence and optical density at 600 nm were measured at different time points during growth in a Multi-detection microplate reader (Synergy, Cytation 3; excitation 325 nm, emission 392 nm)

#### Measurement of FRET-18CSP cleavage in culture supernatants

Supernatants from *S. mutans* UA159 P_*cipB*_luc reporter (SM059) and Δ*htrA* (SM165) were collected by centrifugation (6000 g for 10 min at 4°C) at different time points during growth, corresponding to early, mid-, and late exponential phases. The supernatants were then distributed into the wells of a 96-well plate (115μl in each well), and 5 μl of FRET-18CSP peptide was added (4μM final concentration). The plate was incubated at 37°C in air atmosphere for 1h, and fluorescence was measured in a Multi-detection microplate reader (Synergy, Cytation 3; excitation 325 nm, emission 392 nm).

## RESULTS

### Bioassay for CSP detection in CDM

In this study we used a P*_cipB_*-luc reporter (P_luc_1914) for detection of extracellular CSP activity, as previously described (26), except that high sensitivity was only achieved in the presence of BSA (Figure 2A). BSA was chosen because it is known to enhance competence of *S. mutans* in rich media (37), and in *S. pneumoniae*, BSA prevents CSP proteolysis (38).

**Figure 2.**
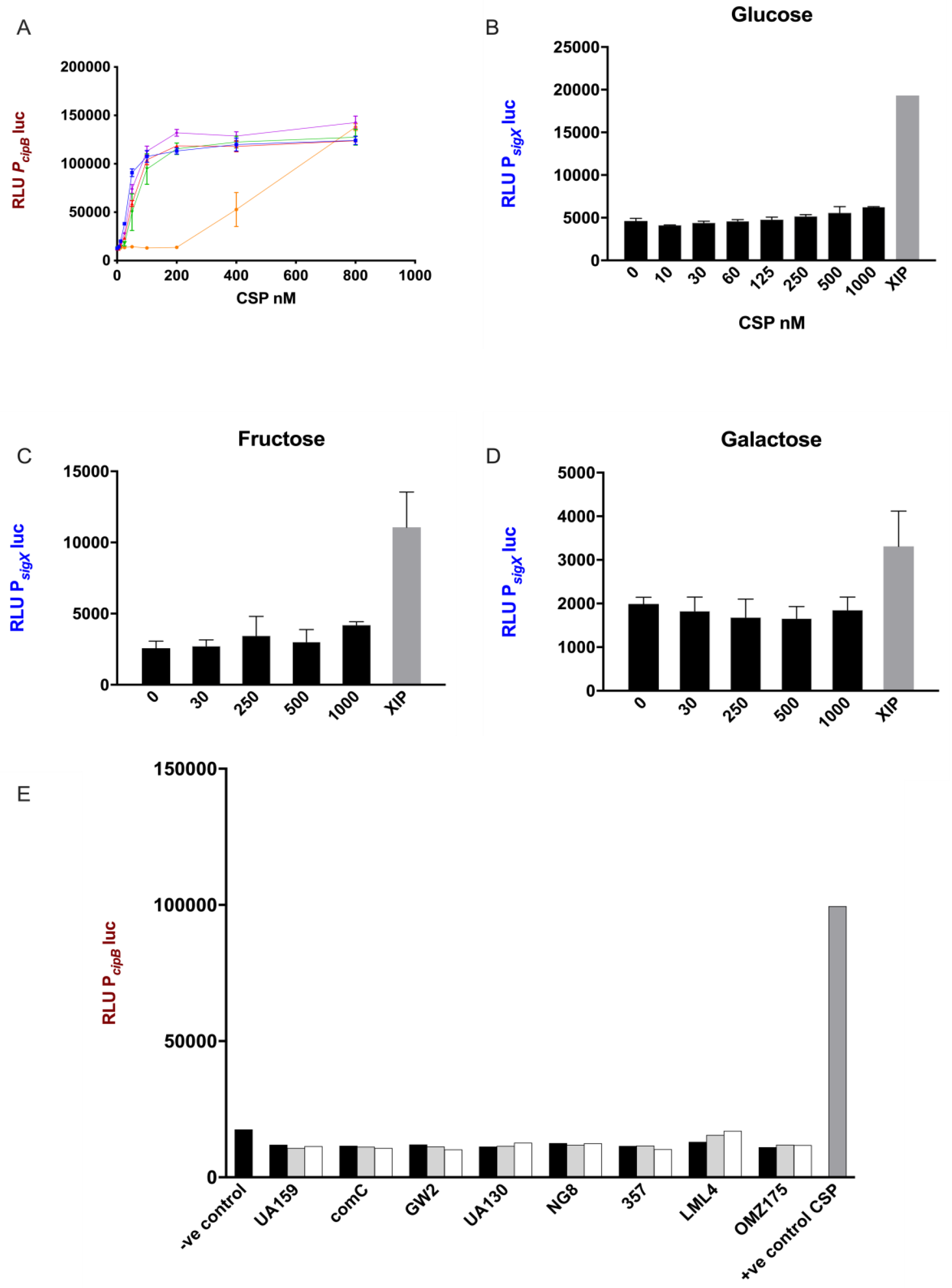
CSP activity in CDM. (A) The indicator strain SM059 (P_*cip*_Bluc) was exposed to serial dilutions of synthetic CSP in the presence and absence of BSA. Different concentrations of BSA were tested. 10% BSA (blue), 5% BSA (red), 1 *%* BSA (green), 0,5% BSA (pink), No BSA (orange). Bars correspond to standard errors of two to three independent experiments; RLU values were measured in a 96 well plate using a Multi-detection microplate reader. (B, C, and D) Activity of P*_sigX_* luc was determined after addition of various concentrations of CSP and 1μM XIP in CDM supplemented with (B) glucose (C) fructose and (D) galactose. Mean and standard deviation for two independent experiments. (E) Supernatants were collected from different *S. mutans* strains at 2 h (black bars), 4 h (gray bars) and 6 h (white bars) growth in CDM and examined for CSP activity using SM059 (P*_cipB_*luc) indicator strain. Medium alone was used as a negative control, and CSP (250nM) was used as a positive control.

The potency of CSP in inducing the activiy of the *cipB* promoter was increased by approximately 33 fold in CDM supplemented with BSA compared with CDM alone (Fig. 2A). BSA concentrations from 0.5% to 10% gave similar results (Fig. 2A).

### Failure of CSP to induce *sigX* in CDM was independent of carbohydrate source

Since the induction of *cipB* was significangly enhanced in the presence of BSA, we decided to further examine the intriguingly fact that CSP fails to stimulate *sigX* in CDM (28–30). Our hypothesis was that increased stimulation of the early response by CSP, as observed with BSA supplementation, would enable the activation of *sigX*, which is a late response in complex media. We used a similar bioassay as described above, but using the promoter of *sigX* linked to the luciferase gene instead. The results showed that CSP concentrations up to 1000 nM failed to induce the P_*sigX*_-luc reporter in CDM supplemented with BSA and glucose (Figure 2B).

Since the carbohydrate source, at least in complex media, has a large influence on the induction of *sigX* expression by CSP (39), we also investigated *sigX* expression in CDM supplemented with fructose and galactose (Figures 2C and 2D). CSP still failed in demonstrating *sigX* induction. Thus, although BSA enables prompt early response to exogenous CSP in CDM, the CSP well-known inducing effect on *sigX* expression in complex media could not be rescued by optimal activation of the early response to CSP.

### Endogenous CSP activity was not detected in the supernatants of *S. mutans*

Lack of endogenous CSP activity in the supernatants of *S. mutans* UA159 grown in CDM has been previously reported by our group (26). It is, however, unknown whether this is a conserved feature in the species. We investigated 6 other strains of *S. mutans*, and the *comC* deletion mutant SM004, for the CSP activity in their supernatants, by growing them in CDM in the presence of BSA (Figure 2E). No *cipB-inducing* activity was detected in culture supernatants collected at early-, mid-, and late-exponential phases of growth, thus indicating that the lack of extracellular CSP activity under such growth conditions may represent a conserved feature in *S. mutans*.

### Suppression of CSP activity during growth in CDM

Since (1) activation of the CSP late response resulting in induction of sigX was abolished in CDM, but not the early response leading to increased expression of *cipB*, and since BSA enhanced the early response, we decided to investigate whether CSP may be degraded over time by *S. mutans*. Supernatants of *S. mutans* cultures exposed to 250 nM synthetic CSP were collected at different time points during growth in CDM. CSP activity in the supernatants of *S. mutans* UA159 was measured by using the *cipB* luciferase reporter described above. Within the first 3 hours, CSP had almost completely disappeared from the culture supernatants (Figure 3A).

**Figure 3.**
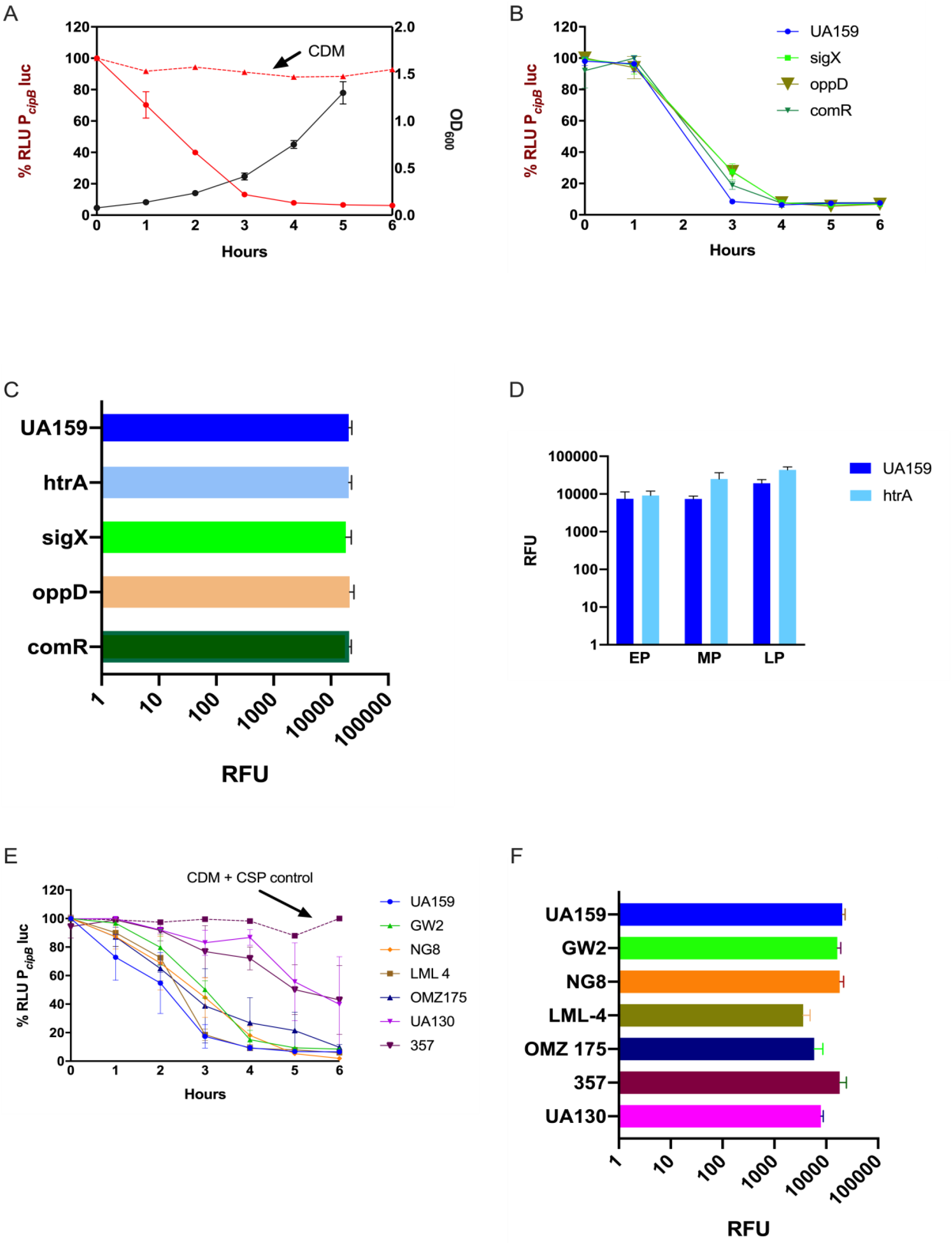
Suppression of CSP activity during growth in CDM. Frozen stock cultures at OD_600_ 0.6 were diluted 10x in CDM containing 10% BSA, and 250nM synthetic CSP (A, B, and E) or 4μM FRET-CSP (C, D, and F) . (A, B, and E) The P*_cipB_*luc reporter was used to measure CSP concentration in the supernatants of different strains (A) *S. mutans* UA159 growth curve was measured by OD_600_ (black line), and indicates the time points when supernatants were collected. CSP activity in CDM alone (positive control) is represented by the red dashed line, and in the supernatants of *S. mutans* UA159, by the red solid line. (B) Loss of CSP activity in the supernatants of UA159, Δ*sigX*, Δ*oppD*, and Δ*comR*. (C and F) CSP cleavage measured at 240 min growth in the presence of the FRET-18CSP peptide, and recorded as relative fluorescence units (RFU) for (C) UA159, Δ*htrA*, Δ*sigX*, Δ*oppD*, and Δ*comR*, *and (F) UA159, GW2, NG, LML-4, OMZ175, 357, and UA130*. (D) FRET-18CSP proteolytic activity in supernatants of UA159 and the Δ*htrA* mutant collected at early (EP), mid- (MP), and late (LP) exponential phase of growth. Error bars show standard error of mean from two to three independent experiments, with 3 parallels each

Due to the recognizable role of CSP in the induction of competence, it was crucial to determine if the loss of CSP activity was dependent on the development of competence. For this, we investigated the CSP activity in the culture supernatants of UA159 deletion mutants for *sigX, oppD*, and *comR* (Table). In all of them, a dramatic reduction in activity of exogenously added CSP was observed (Figure 3B).

To determine whether such an inhibitory effect on CSP activity could be due to proteolytic cleavage, we supplemented the CDM medium with a FRET-18CSP reporter peptide (Figure 3C). We found that all three isogenic mutants (*sigX*, *oppD*, and *comR*) grown under such conditions promoted increase in fluorescence activity, as measured after 240 min incubation at 37°C (Figure 3C), thus suggesting that the ability of *S. mutans* to proteolyticially inactivate CSP does not require expression of key elements of the competence regulon. We also examined the effect of inactivating the *htrA* gene coding for the *S. mutans* HtrA protease, which in *S. pneumoniae* inactivates the CSP and cleaves misfolded proteins. No reduction in CSP degradation was observed for the *htrA* mutant (Figure 3C). CSP activity was indeed slightly higher for the culture supernatants of the *htrA* mutant at mid- and late-exponential phases of growth (Figure 3D).

Finally, we tested whether suppression of CSP activity is extended to other *S. mutans* strains. In all the strains tested, reduction in CSP activity was observed, though at different levels (Figure 3E). When grown in the presence of the FRET-18CSP reporter peptide, an increase in fluorescence was observed for all strains. We conclude that *S. mutans* suppresses CSP activity during growth in CDM, and that the mechanisms involved are active in the absence of *htrA, sigX, comR* or *oppD*, indicating the involvement of competence-regulated independent factors. Proteolytic activity leading to CSP inactivation seems to be conserved in *S. mutans*.

### XIP activity is detected in the culture supernatants of practically all *S. mutans* strains grown in CDM

*S. mutans* UA159 produces and responds to exogenous XIP in CDM (4, 8). We investigated whether XIP production is conserved in *S. mutans*. We used the P*_sigX_*-luc Δ*comS* indicator (SM091) to estimate the XIP concentration in the supernatants of the different strains during growth. The minimum detection level was set at 10 nM, as determined by standard curve measurements using synthetic XIP. Our results showed the presence of extracellular XIP activity in all strains tested. The highest XIP concentrations in the supernatants of growing cultures ranged from approximately 200 to 26 000 nM (Figure 4A), with most showing XIP accumulation starting at the mid-exponential growth phase, and reaching maximal values at early stationary phase (data not shown). We conclude that the growth conditions that support extracellular XIP pheromone accumulation by UA159 may also sustain XIP production by a majority of the strains.

**Figure 4.**
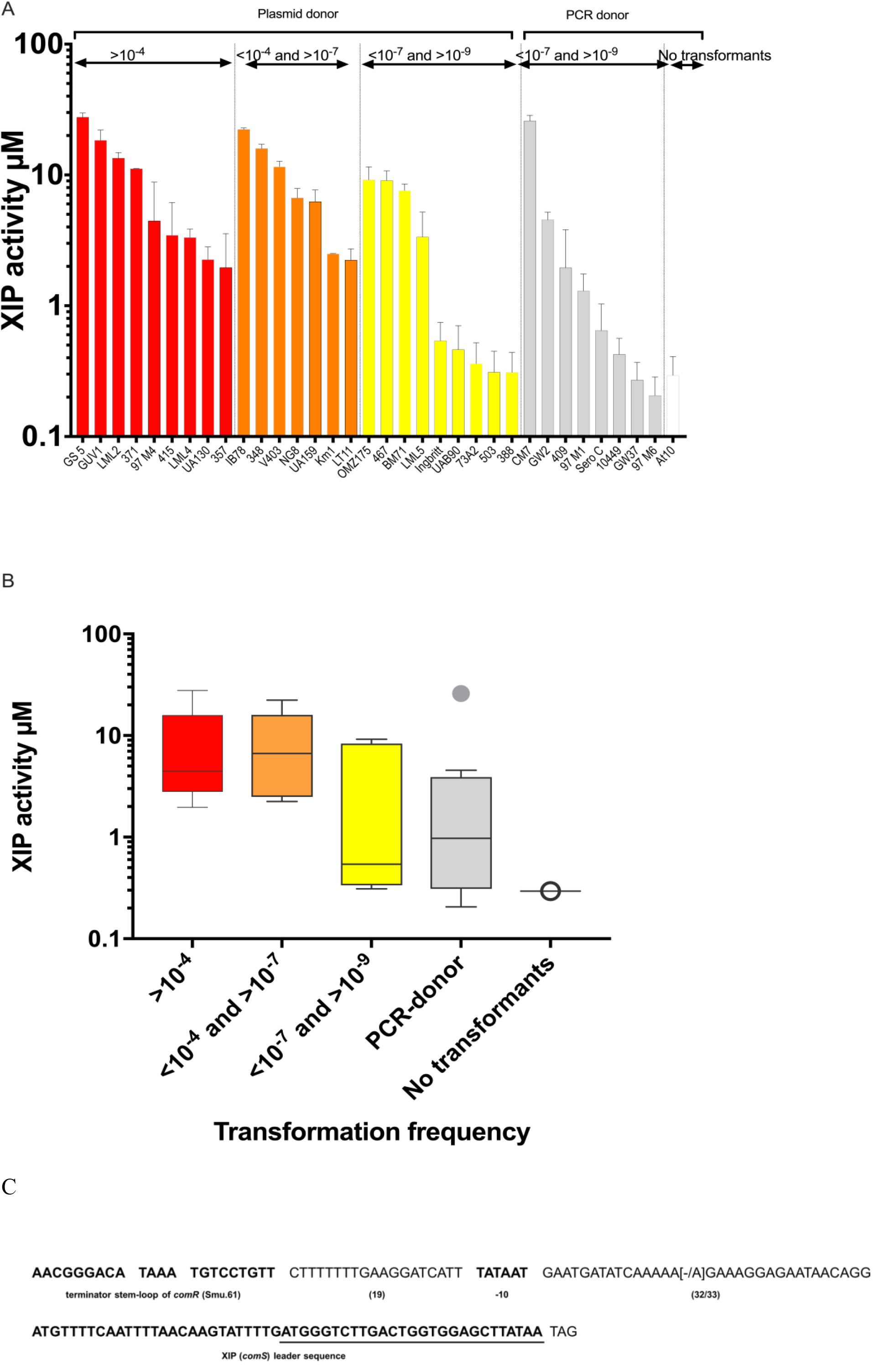
Conservation of XIP production and transformation in CDM. (A) Extracellular XIP concentration was measured in the supernatants collected from *S. mutans* strains grown overnight at 37°C in air. (a) Columns show average of XIP-equivalent concentration from 3 independent experiments. Bars correspond to standard error. The indicator strain used was a Δ*comS* P*_sigX_*-luc reporter (SM091). Different colors indicate transformation frequency in CDM supplemented with synthetic XIP by a plasmid donor (pVA838): (>10^−4^ in red, between 10^−4^ and 10^−7^ in orange, and between 10^−7^ and 10^−8^ in yellow). For those that were not transformed by the plasmid donor, transformation frequency with a 6.3 Kb PCR donor designed to integrated into the chromosome by homologous recombination are shown (all frequencies below 10^−7^; grey). Only At10 yielded no transformants (white). (b) Whisker plots showing the average of XIP activity of the strains shown in (a), grouped according to transformation frequency. (C) Conservation of *comS* promoter and *comS* sequences in *S. mutans* strains. The first sequence in bold shows the terminator stem-loop of *comR* (Smu.61) thought to function as part of the *comS* promoter. Downstream the stem-loop is a 19 bp sequence, followed by the putative −10 element of the *comS* promoter distant 32 to 33 bases from the *comS* gene sequence shown in bold. Underlined is the sequence corresponding to the XIP functional heptapeptide (ComS_11-17_ – GLDWWSL). Variable bases compared with *S. mutans* UA159 are in brackets.

### Thirty-three out of thirty-four strains were naturally transformable

Natural transformation without the addition of synthetic pheromone was detected in 23 of the 34 strains (data not shown). Because optimal levels of XIP may not be present at the time of competence under *in vitro* conditions, it was of interest to learn whether addition of synthetic XIP would result in higher transformation levels. Our results showed that synthetic XIP extended the range of transformed strains. Three of the strains with detectable XIP levels in the supernatants, including OMZ175, UAB90, and KM1 were transformed only upon addition of synthetic XIP. In the strains that were not transformed with the initial protocol, we increased the time during which they grew in the presence of synthetic XIP and donor DNA to 10 h, but even then no transformants were obtained. (40). We next used a more sensitive protocol based on the use of PCR large fragments as donor DNA (35, 36). Only strain At10 was not transformed, thus indicating that a majority of *S. mutans* are amenable to transformation. Overall, strains that exhibited lower transformation frequencies showed lower levels of XIP activity in their culture supernatants (Figure 4B).

### The *comS* gene was conserved in the 34 *S. mutans* strains

The *S. mutans comS* gene encoding the XIP precursor is essential for competence development. Given the variability in transformation efficiency observed above, and that some strains were not amenable to transformation, we investigated whether the *comS* gene and its putative promoter sequences were conserved in the *S. mutans* strains included in the study. The results showed that *comS* was identical in all the 34 strains (Figure 4C). Highly conserved sequences were also found in the *comS* putative promoter. In this region the only difference in relation to UA159 was the presence or absence of an additional adenine between the −10 element and the *comS* translation initiation site. The additional adenine was present in 22 out of the 34 strains evaluated. In all the strains, *comS* was located 57-58 nt downstream of the tRNA transcriptional terminator of *comR* (SMU_61) and the putative −10 element was 32-33 nt upstream of the *comS* translation initiation site. While this study was being performed, new genomic sequences for more 78 strains of *S. mutans* were made available on public databases (41). In order to compare our results with the new genomic sequences we ran BLAST using the sequence shown in Fig. 5 against the CoGe database (http://genomevolution.org/CoGe/CoGeBlast.pl) (42). The *comS* sequences in these strains were identical to the *S. mutans* UA159. In the *comS* promoter the sequences varied in only a single base pair at the same position as that observed in 22 of the strains sequenced in our study. Taken together the results indicate that the *comS* gene and promoter regions are highly conserved in the *S. mutans* strains analyzed, suggesting that variations in transformation levels and XIP production are not correlated with differences in this region.

## DISCUSSION

In complex media, the CSP pheromone triggers the expression of bacteriocin-related genes, followed by a late response characterized by increased expression of the XIP-encoding gene *comS*, and competence development (43, 44). The mechanisms leading to activation of *comS* expression and competence remain unknown. It is also still unknown the reason why CSP fails in stimulating competence in peptide-free defined medium. It has been suggested that *S. mutans* may perhaps produce a protease that, similar to the HtrA protease in *S. pneumoniae*, could inactivate the CSP (29). This possible explanation, however, did not seem to match with the later finding that CSP is actually active in CDM, in that it can stimulate *cipB* and all other genes regulated by ComED (28). Our results show, however, that *S. mutans* can indeed inactivate CSP by a mechanism that most probably involve proteolysis, given the results obtained with the FRET-18CSP reporter peptide. We confirmed previous results that CSP induces cipB (28), and found that a CSP concentration as low as 1 nM in the presence of BSA was sufficient to activate the early bacteriocin response. This is in oppose to the CSP concentration required to activate *sigX*, which can be observed with values that exceed the maximum concentration used in this study (1000 nM) (data not shown). Inactivation of the *S. mutans* CSP by proteases produced by other streptococcal specieshave been known for more than ten years (45). However, this is the first indication that *S. mutans* can also inhibit the activity of its own CSP, most probably via proteolytic activity. Unlike for *S. pneumoniae* (38), however, the HtrA protease in *S. mutans* was not found to be required for CSP inactivation.

In *S.pneumoniae*, luciferase and gfp reporter strains for CSP activity have been successfully used to identify S. pneumoniae CSP pheromones in culture supernatants (46). Also in the seminal study that identified the *S. pneumoniae* competence factor, the CSP was isolated and purified from the supernatant (47). Also for *S. mutans* UA159 grown in complex medium, CSP has been identified by mass spectrometry in the supernatant fraction (23). Based on these findings, we hypothesized that if *S. mutans* produces endogenous CSP, we would probably be able to detect it in its culture supernatant. The protocol we developed in this study using a *cipB* luciferase reporter suspended in CDM supplemented with BSA enabled the identification of concentrations as low as 1 nM. Despite this high sensitivity, we could not detect any CSP inducing activity in the supernatants of the different strains tested, indicating therefore that endogenous CSP was either absent, present at very low levels or not present in the supernatant fraction.

In contrast to CSP, endogenous XIP activity was present in the supernatants of most strains. Moreover, activation of the XIP system in CDM enabled transformation of all but one of the 34 strains tested. The *comS* gene encoding the pre-processed form of XIP was indeed found in all the strains, which is in line with the results of a recent study reporting *comS* as part of the *S. mutans* core genome (11). In general, higher levels of XIP detection in the supernatants correlated with increased transformability. Of note, some strains that produced high concentrations of endogenous XIP, needed to be stimulated by synthetic XIP to transform. This was, however, not surprising, given the fact that maximum concentrations were in most cases observed only close to the stationary phase, which is a period during which competence under laboratory conditions is already shut-off. For some of the strains that produced low levels of XIP, transformation was only detected by using a large PCR amplicon as DNA donor, which is known to result in increased recovery of transformants (35, 48).

Taken together, our results indicate that the XIP system is conserved in *S. mutans*, and that all *S. mutans* strains may belong to a single pherotype. Moreover, the conserved ability of *S. mutans* to cleave CSP under conditions that favor accumulation of XIP may provide an adaptive advantage to *S. mutans*, by allowing them to fine tune competence and bacteriocin responses in response to environmental changes. While the knowledge on pheromone signaling for any species is mostly based on models derived from limited selected strains, understanding how the system works in different strains of a species is of high relevance for the development of any new strategies aiming at interfering with their communication systems.

## ACKNOWLEDGMENTS

We thank Andreas Podbielski for the kind gift of pFW5-luc and Roy Russel for providing the *S. mutans* strains LML-2, LML-4, LML-5 and At10. We are grateful to Anne Karin Kristoffersen for helpful assistance in DNA sequence analysis. This study was financed in part by the Coordenação de Aperfeiçoamento de Pessoal de Nível Superior - Brasil (CAPES) - Finance Code 001 (A.P.R.F. postdoctoral grant), and by the program International Partnership for Outstanding Education, Research, and Innovation (INTPART), RCN grant number 274867.

